# Mammalian oocytes receive maternal-effect RNAs from granulosa cells

**DOI:** 10.1101/2025.02.10.637575

**Authors:** Caroline A. Doherty, Abdulfatai Tijjani, Steven C. Munger, Diana J. Laird

## Abstract

It is currently thought that growing mammalian oocytes receive only small molecules via gap junctions from surrounding support cells, the granulosa cells. From the study of chimeric preantral oocyte and granulosa cell reaggregations, we provide evidence that growing mouse oocytes receive mRNAs from granulosa cells. Among the >1,000 granulosa-transcribed RNAs we identified in the oocyte, those that contribute to proper oocyte maturation and early embryo development were highly enriched. Predicted motifs for two RNA-binding proteins that function in RNA trafficking, FMRP and TDP43, were abundant in the UTRs of the granulosa-derived transcripts. Immunostaining demonstrated that both FMRP and TDP43 co-localize with the actin-rich granulosa cell protrusions that span the zone pellucida and connect to the oocyte, suggesting their role in importing mRNAs. Our results offer the possibility that oocyte failure may not always reflect an intrinsic oocyte deficiency but could arise from insufficient supply of maternal transcripts by granulosa cells during oocyte growth.

## Introduction

Mammals are born with a finite supply of immature, quiescent oocytes that are gradually recruited for growth over time. Mouse oocytes stockpile biosynthetic components as they increase in volume by approximately 300-fold^1^. Many of these components are general metabolites while others are transcripts and the corresponding protein products that support the oocyte-to-embryo transition^2^. These RNAs and proteins in the oocyte are produced by maternal effect genes^3,4^. Identifying the minimal components necessary for proper oocyte maturation and the mechanisms by which oocytes accumulate and organize these components could offer new solutions to improving fertility outcomes. How oocytes grow so large, and on what timescales, differs across species. Yet one common feature is conserved – the transfer of biosynthetic products to oocytes from neighboring support cells^5^.

Following their differentiation in the fetal ovary, immature mouse oocytes become surrounded by a squamous layer of epithelial cells called granulosa cells. The unit of an oocyte enclosed in somatic cells is called a follicle. As subsets of quiescent, primordial follicles are recruited to grow, the oocyte begins secreting a layer of glycoproteins called the zona pellucida, which separates the oocyte from the granulosa cells^6^. As the follicle grows, the granulosa cells become cuboidal (primary follicle) and then proliferate to become multilayered (secondary follicle; **Fig 1A**). Despite being physically separated from the oocyte by the zona pellucida, granulosa cells grow thin filopodia-like projections, called transzonal projections (TZPs), that penetrate the glycoprotein matrix and contact the oocyte (**Fig 1B**). TZPs are conserved across mammals, from mice to humans (**SFig 1A,B**). The growing follicle develops a fluid-filled cavity called an antrum (antral follicle) and is eventually ovulated^5,7,8^ (**Fig 1A**).

**Figure 1.**
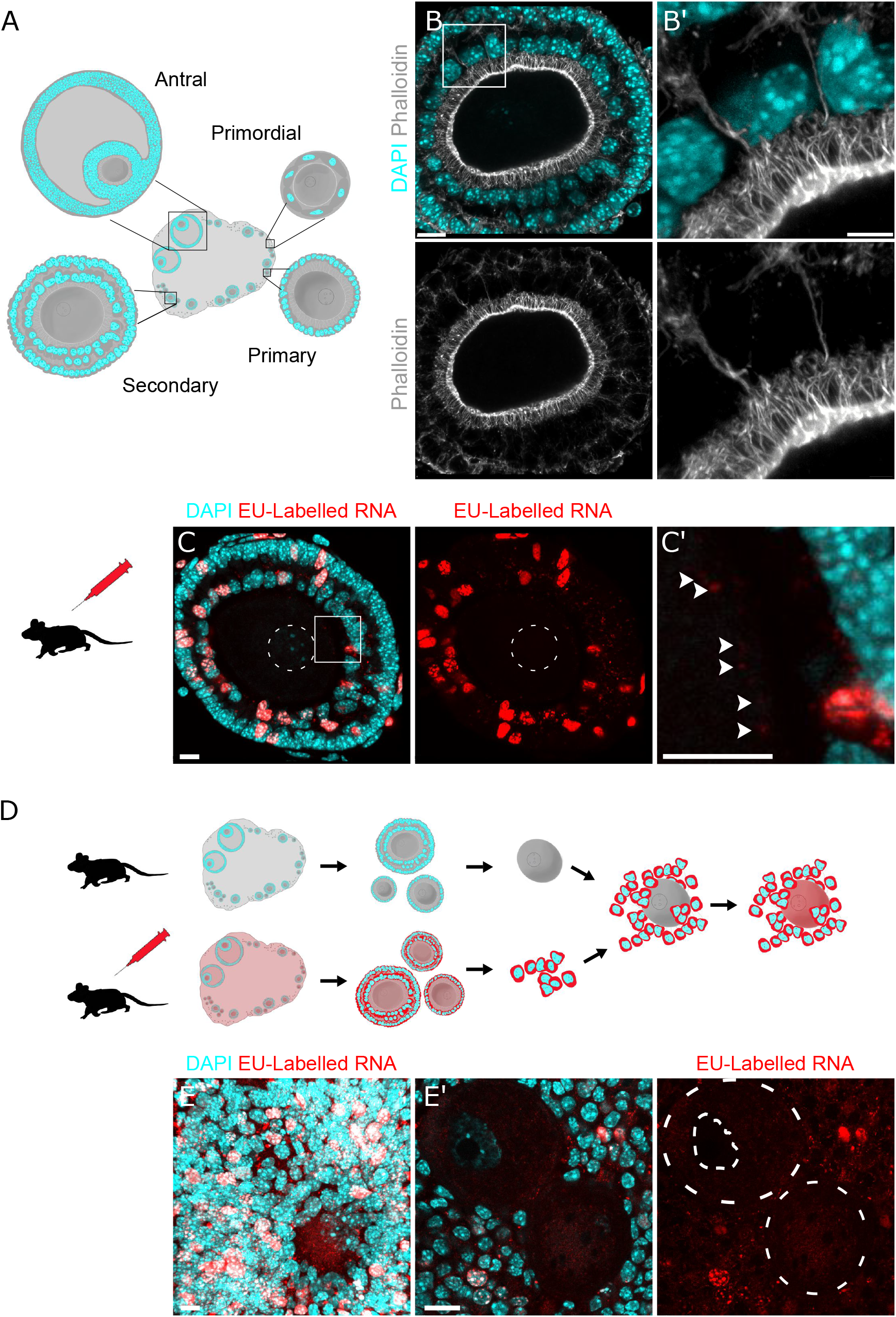
Labeled RNA in granulosa cells can be detected in preantral oocytes. A. Schematic of folliculogenesis: *Primordial follicles* are quiescent oocytes present from birth that are surrounded by a layer of squamous granulosa cells. *Primary follicles* are growing follicles that have begun secreting a layer of glycoprotein, the zona pellucida, that separates the oocyte from the granulosa cells. To maintain contact with oocytes across the zona pellucida, granulosa cells grow thin, filopodia-like projections, called transzonal projections (TZPs). *Secondary follicles* have multiple layers of granulosa cells. Both inner and outer granulosa cells grow TZPs that contact the oocyte. *Antral follicles* grow a large fluid-filled sac and are the final stage of oogenesis prior to ovulation. B. NSPARC z-projection of an intact mouse multilayered secondary follicle. Granulosa cell nuclei marked with DAPI in cyan and transzonal projections (TZPs) marked with phalloidin in grayscale. A magnification (B’) highlights a TZPs contacting the oocyte from both the inner and outer layers of granulosa cells. Scale bars = 10 um. C. Wholemount confocal z-projection of a secondary follicles processed with Click-iT kit 16 hours after ethynyl uridine IP injection. DNA labelled with DAPI in cyan and nascent RNA labelled with EU in red. Nascent RNA does not appear in the oocyte nucleus (outlined in white dashes) however it does appear in small puncta at the oocyte cortex seen in the magnification (C’) noted with arrowheads. Scale bars = 5 um. D. Re-aggregation scheme whereby unlabeled oocytes are re-aggregated with granulosa cells labeled with EU. The granulosa cells were isolated 16 hours after IP injection of EU. After six days of culture the aggregates are fixed and processed with a Click-iT kit to visualize EU-labeled RNA. E. Wholemount z-projection of reaggregates and single confocal slice (E’). EU-labelled puncta appear in the oocytes, indicating transfer of RNA from granulosa cells to oocytes. Scale bars = 10 um.

It is currently thought that oocyte support from granulosa cells is limited to small molecules, like metabolic precursors, conveyed via gap junction connections at the tips of TZPs. Therefore, it is presumed that all macromolecules in the oocyte cytoplasm larger than 1 kDa– the upper size limit of a gap junction pore^9^ were synthesized by the oocyte itself. For example, GFP protein is approximately 27 kDa and GFP mRNA is approximately 234 kDa, both too large to fit through a gap junction^10,11^. However, demonstration that GFP-tagged proteins expressed in cultured bovine granulosa cells can be detected in un-transfected oocytes following co-culture raises the possibility that larger transcripts or protein can be transferred to the bovine oocyte^12,13^.

In *Drosophila*, oocytes rely on support cells to synthesize many of their components and maternal effect transcripts. Because mammalian oocytes are similarly paused in the diplotene stage of prophase I from primordial stages until nuclear envelope breakdown ^14,16^, we investigated whether mouse oocytes also receive transcripts from support cells.

## Results

### Granulosa cells re-establish contact with oocytes following enzymatic separation and reaggregation

Previous studies of granulosa cell-oocyte interactions in the transzonal region were performed on cumulus oocyte complexes removed from antral follicles^15^. However, the rate of oocyte growth plateaus at antral stages^17,18^ (**SFig. 1C**), suggesting that biosynthetic requirements peak earlier in growth during primary and secondary stages. Therefore, we developed *ex vivo* protocols for investigating the dynamics of granulosa cell-oocyte interactions during pre-antral stages. Previous work demonstrated that mechanically separated granulosa cells re-grow TZPs and re-attach to oocytes isolated from antral follicles^15^. To scale up our experimental pipeline we isolated granulosa cells and oocytes from primary and secondary stage follicles of juvenile mouse ovaries by enzymatic digestion followed by filtering and then reconstructed preantral follicles *in vitro* by reaggregating granulosa cells with oocytes (**SFig 1D,E**). Imaging with phalloidin confirmed that granulosa cells regrow oocyte-directed TZPs in these follicle reaggregates (**SFig 1F**). A similar enzymatic separation and reaggregation protocol with longer culturing timelines gave rise to live pups^19^.

### Labelled RNA transits from granulosa cells to oocytes

To investigate whether RNA can transit from granulosa cells to the oocyte, we labeled newly synthesized RNA *in vivo* and then analyzed intact and reaggregated preantral follicles. We injected postnatal day 12-15 mice with the uridine analog 5-ethylnyl-uridine (EU), which incorporates into nascent RNA, isolated the pre-antral follicles 16 hours later, and detected the EU with Click Chemistry. In wholemounted follicles we observed variable labeling of granulosa cells by EU, likely due to their proliferation at this stage. We did not detect nascent RNA in the oocyte nucleus; however, we did observe EU-labeled RNA puncta present at the oocyte cortex (**Fig 1C**). To exclude the possibility that EU-RNA was synthesized in oocytes, we reaggregated granulosa cells isolated from EU-injected mice with oocytes from an unlabeled mouse. After growth of this reaggregated follicle for six days *in vitro*, EU-labeled RNA was detected in the oocytes (**Fig 1D,E**). This result indicates that RNA can transit from mouse granulosa cells to oocytes.

### Chimeric follicle reaggregates identify transferred transcripts that are involved in oocyte maturation and preimplantation development

To identify specific RNAs that move from the granulosa cells to the oocyte during pre-antral stages, reaggregation was carried out with granulosa cells and oocytes from two strains of inbred mice, C57BL/6J (B6) and CAST/EiJ (CAST). These distantly related strains differ at nearly 18 million single nucleotide polymorphisms (SNPs) across the genome, with over 400,000 variants identified in annotated transcripts^19^. This high degree of transcriptomic diversity enables the identification of strain origin for many expressed genes, as nearly half of 100bp RNA-sequencing (RNA-seq) reads would be expected to harbor one or more SNPs that differ between these strains. After reaggregating B6 oocytes with CAST granulosa cells, we re-isolated and purified the B6 oocytes, and then performed bulk RNA-seq on three biological replicates of pooled oocytes (**Fig 2A**). Following a stringent alignment of reads to strain-specific transcriptomes, we identified approximately 1,400 transcripts present in all three B6 oocyte replicates that most likely originated from the CAST granulosa cells (**Supplementary Table 1**). Of the 500 transcripts with highest read counts, 273 were present in all three independent replicates, giving us high confidence in the reproducibility of our findings (**Fig 2B**).

**Figure 2.**
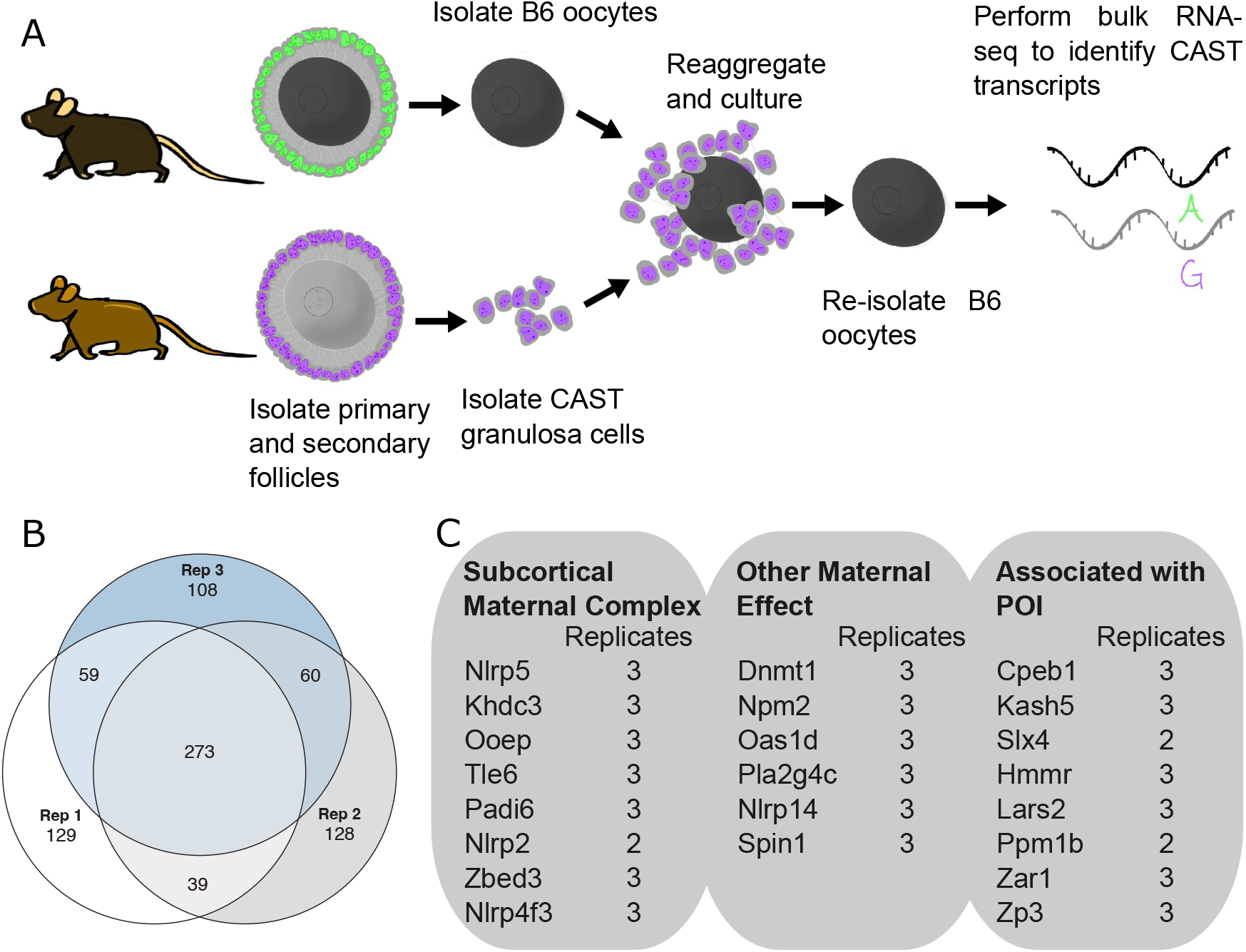
Identification of granulosa-derived transcripts in preantral oocytes. A. Schematic of follicle re-aaggregation protocol combining primary or secondary stage B6 oocytes (black) with Cast granulosa cells (purple). B. Out of the top 500 genes with CAST SNPs with most abundant reads identified in RNA-seq from B6 oocytes, 273 were present in all three follicle reaggregation replicates (Rep). C. Relevant transcripts identified that are associated with the subcortical maternal complex, maternal-effect genes, and genes associated with primary ovarian insufficiency.

Next we performed pathway enrichment analysis and functional enrichment analysis on the 1,336 protein-coding genes from CAST present in all three B6 oocyte replicates^20,21^. The Mammalian Phenotype Ontology identified abnormal oogenesis (MP:0001931), abnormal female germ cell morphology (MP:0006361), and maternal effect (MP:0003718) in the top 5 most enriched categories^22^. The pathway with the largest enrichment effect from the Kyoto Encyclopedia of Genes and Genomes (KEGG) database was pyruvate metabolism (mmu00620)^23^. The Subcortical Maternal Complex (SCMC), which is required for storage of maternally-deposited RNAs and proper oocyte maturation and embryogenesis, was the cellular component with the largest enrichment identified by Gene Ontology (GO:0106333) ^24–27^. Specifically, seven of the eight genes encoding components of the SCMC were present in all three of our replicates and the eighth was present in two replicates (**Fig 2C**). Other maternal effect genes that may be associated with the SCMC, but also may have other functions, were also present in all three of our replicates. Additionally, transcripts for the maternal-effect gene *Zar1* were present in all three replicates. Mutations in *Zar1* are also associated with primary ovarian insufficiency (POI), a polygenic condition resulting in a premature exhaustion of the oocyte pool. Several other transcripts for genes associated with POI ^28,29^ were also present in all three of our replicates (**Fig 2C**). Together these analyses reveal that granulosa cell-derived transcripts in the oocyte function in oocyte maturation and early embryo development.

### Trafficking-associated RNA-binding proteins co-localize with TZPs

RNA-binding proteins (RBPs) perform diverse functions in the outcome of an RNA molecule and are involved in translation regulation, RNA stability, and trafficking among other roles. When RNAs are trafficked, they associate in phase-separated structures with RNA-binding proteins called ribonucleoprotein particles (RNPs)^30^. RNPs, though small, can be resolved with light microscopy. If RNAs are moved from granulosa cells to oocytes they would likely be packaged in RNPs.

We used a RBP motif analysis resource that ranks RBPs based on the number of motif instances present in our list of transferred RNAs^31^. Out of the top 50 RBPs with the most motif instances in our collection of RNAs, both TDP-43 and FMRP stood out because they are well-characterized for their roles in RNA-trafficking in neurons^32,33^ (**Supplementary Table 2**). Additionally, FMRP is well-known to be associated with Primary Ovarian Insufficiency and FMRP-GFP transfected into bovine granulosa cells has been observed to transit to oocytes^12,13,34^.

We investigated whether TDP-43 and FMRP participate in RNA transfer within follicles. Using confocal microscopy we detected colocalization of TDP-43 and FMRP puncta with TZPs in whole-mount preantral follicles (**Fig 3A,B**). To further characterize FMRP localization within the follicle we used super-resolution microscopy and observed what appear to be TZP-associated FMRP puncta docked at the oocyte membrane (**Fig 3A**). These results are consistent with the hypothesis that TDP-43 and FMRP are involved in transferring RNAs via TZPs.

**Figure 3.**
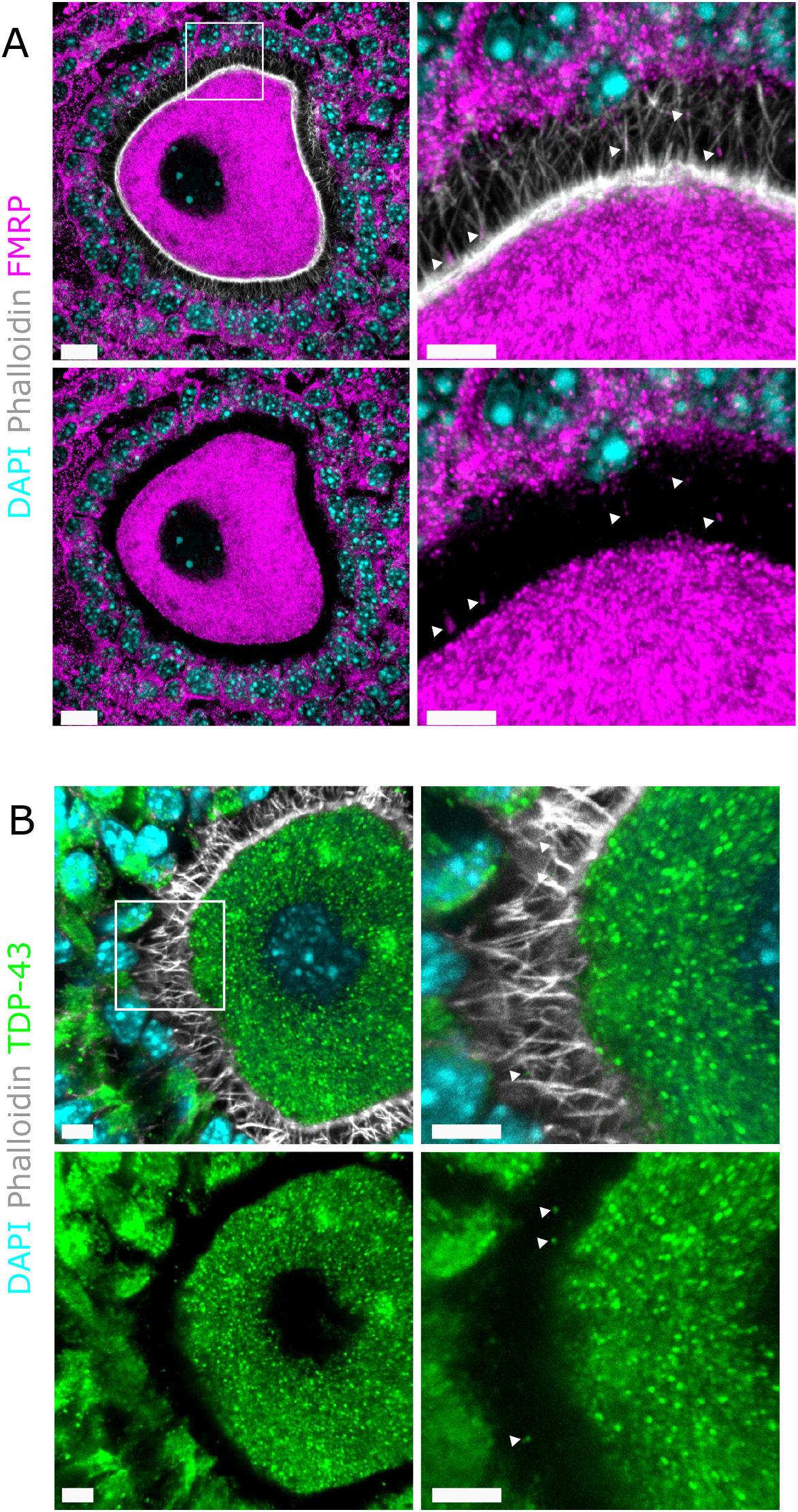
RNA-Binding Proteins predicted to bind to trafficked RNAs localize in TZPs. A. Immunostaining and super-resolution microscopy demonstrate co-localization of FMRP (pink) with TZPs (stained with phalloidin in grey; see arrowheads) and “docking” of FMRP with the oocyte cortex. DAPI marks nuclei in turquoise. Scale bars = 10 um. B. TDP-43 (green) similarly localizes with TZPs (grey; see arrowheads). Scale bars = 5 um.

### Vesicles co-localize with TZPs

Lipid bilayers generally do not allow for the passage of large, charged molecules such as RNA^35^. However, RNA could cross the granulosa and oocyte cell membranes if packaged in vesicles. To see whether vesicles play a role in granulosa to oocyte trafficking we imaged follicles expressing the membrane-marking reporter *Rosa26*^*mTmG*^. The modified MARCKS tag linked to tdTomato causes the fluor to associate with hydrophobic bilayers and therefore also labels vesicles ^36–38^. Upon imaging of the tdTomato signal, we observed punctate structures that co-localize with TZPs (**Fig. 4A**). This suggests that vesicle trafficking occurs in TZPs.

**Figure 4.**
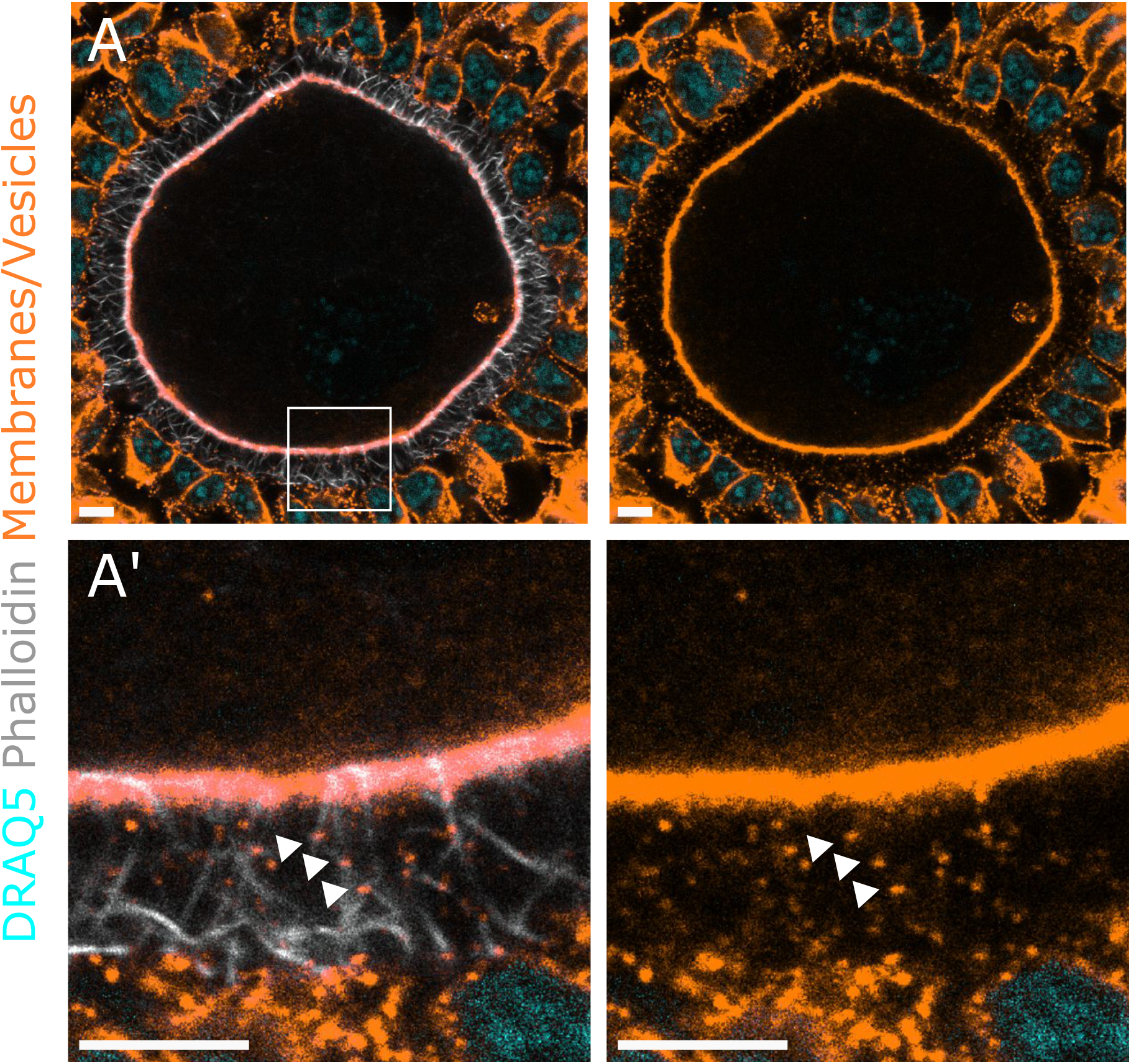
Vesicles co-localize with TZPs. A. Wholemount confocal slice and magnification (A’) of a mouse follicle expressing the mTmG reporter, labeling membranes and vesicles with tdTomato in orange. TZPs labeled with phalloidin in grayscale and DNA labelled with DRAQ5 in cyan. Magnified image demonstrates co-localization of vesicles with TZPs. Scale bars = 5 um.

## Discussion

Oocytes accumulate stockpiles of maternally derived transcripts that must be stored until the oocyte to embryo transition. This accumulation process was previously thought to be oocyte-intrinsic in mammals. Taking advantage of follicle reaggregation assays and strain-specific transcriptomic analysis, we present evidence that mouse oocytes receive RNAs, including transcripts from known maternal-effect genes, from the somatic granulosa cells.

The chance of achieving a successful pregnancy without intervention decreases with maternal age. This age effect has been attributed to the oocyte rather than the uterus since the success rate of embryos from young donors does not vary by recipient age^39^. While evidence points to accumulated damage and breakdown of proteins that hold meiotically arrested chromosomes together as causes for declining oocyte quality^40,41^, emerging work on the ovarian microenvironment suggests that oocyte health may also be controlled by the mechanical and metabolic properties of the surrounding ovarian tissue^42,43^. Consistent with this idea, other studies find that an age-dependent decrease in the density of TZPs that extend from granulosa cells and that old oocytes can be rejuvenated by younger granulosa cells^43,44^. Our findings raise the possibility that as ovaries age, granulosa cells become less effective at transferring important RNAs to oocytes and therefore older oocytes are insufficiently “loaded.”

The function of the the granulosa-derived transcripts we identified in the oocyte raises the possibility that the transfer process promotes health of the oocyte and embryo. One salient category is maternal effect genes, including *Zar1*, which promotes coalescence of oocyte structures known as mitochondrial-associated membraneless compartments that store maternal mRNAs necessary for oocyte maturation^45^. The *Zar1* mechanism resembles that of the Drosophila gene *oskar. Oskar* mRNA is synthesized in the supporting nurse cells of the fly ovary and is then actively localized to the oocyte posterior where its protein product nucleates the formation of the germ plasm, a collection of stored mRNAs necessary for germline specification^46,47^. In mice, *Zar1*-null oocytes grow to their full volume and can be ovulated, although embryos subsequently arrest at the 1 or 2-cell stage. Our identification of a specific set of transcripts for oocyte maturation, fertilization and embryo development that are transcribed in granulosa cells and conveyed to the growing oocyte suggests an evolutionarily conserved reliance of oocytes on specific macromolecular support from somatic cells.

Our studies provide evidence that RNA transfer occurs with the stabilization of RBPs within extracellular vesicles that travel along TZPs. TZPs act as cellular bridges that allow for the transport of RNP complexes from granulosa cells to the oocyte. *FMR1* pre-mutations are well-known to be associated with infertility^50,52^ yet, how these mutations compromise infertility mechanistically is unknown. We provide evidence that FMRP and TDP-43 may be involved in transferring RNAs to the oocyte via TZPs. In specialized cellular structures, such as filopodia-like tunneling nanotubes and cytonemes, there is evidence that the actomyosin network traffics vesicles over long distances^54^. TZPs share many physical and morphological similarities with these structures^7^. Therefore, vesicle-mediated transfer of granulosa-derived RNAs is consistent as a mechanistic capability of TZPs. This is supported by the presence of vesicles in TZPs. The mechanism of transport as well as the cargo both constitute important future targets for assisted reproductive therapies.

## Methods

### Mouse handling

All mouse work was performed under the University of California, San Francisco (UCSF) Institutional Animal Care and Use Committee guidelines in an approved facility of the Association for Assessment and Accreditation of Laboratory Animal Care International.

### Follicle isolation and staining

Follicles were mechanically isolated in warmed L15 media (11415064 Gibco) on a heated stage by ripping ovaries with forceps. Enzymatic isolation of follicles with collagenase or Liberase causes TZPs to collapse. Follicles were transferred via EZ-Grip pipette to fresh/warm L15 media and then fixed in 2% PFA (cat 043368.9M Thermo Scientific) for 10 minutes with the PFA being directly added to the media. All wash/incubation sets were performed in glass-well dishes on an orbital shaker. Follicles were then washed 3 times for 5 minutes in 0.05% PBS-Tween (cat P7949-500ML Sigma-Aldrich) and then permeabilized in 0.01% PBS-Triton (cat X100-1L Sigma-Aldrich) for 20 minutes or one hour specifically for the anti-mCherry/tdTomato staining. Blocking was performed in 0.05% PBS-Tween with 5% donkey serum (S30-100mL Sigma-Aldrich) and 1% BSA (cat A30075 RPI Research Products) and then incubated overnight with primary antibodies diluted in Toyobo Can Get Signal Solution A (cat NKB-501 Toyobo). Follicles were then washed 3 times for 5 minutes in 0.05% PBS-Tween and then incubated for 2 hours in secondary antibody diluted in Toyobo Can Get Signal Solution B (cat NKB-601 Toyobo).

Some samples were then incubated with WGA for 1 hour, Phalloidin for 30 minutes, DAPI for 15 minutes or DRAQ5 for 15 minutes. When staining was finished, follicles were mounted in a 50/50 mixture of Aqua Polymount (18606-20 Polysciences) and RapiClear 1.52 (RC1520001 SUNJin labs) on slides with coverslips for imaging.

List of stains and primary/secondary antibodies used:

Phalloidin 405 (A30104 Invitrogen) or 488 (A12379 Invitrogen)
DAPI (EN62248 Invitrogen), DRAQ5 (62251 Thermo Scientific)
WGA 680 (W32465 Invitrogen)
Goat anti-FMRP (Knockdown validated, PA5-18742 Invitrogen)
Rabbit anti-TDP43 (Knockout validated, AB109535 abcam)
Rabbit anti-mCherry (Cross-reacts with tdTomato, AB167453 abcam)
Donkey anti-goat 488 (A11055 Invitrogen), 594 (A11058 Invitrogen)
Donkey anti-rabbit 488 (A21206 Invitrogen), 555 (A31572 Invitrogen), 594 (A21207 Invitrogen)

### Baboon and human tissue acquisition

Baboon ovaries were obtained from the Southwest National Primate Research Center biomaterials program. Deidentified human ovaries were obtained from Donor Network West through the UCSF Vital Core. Organs were transported on ice in Belzer UW Cold Storage solution (Preservation Solutions, Inc. Elkhorn, WI). Baboon and human samples were processed with a combination of a tissue grinder kit (CD1-1KT Sigma-Aldrich) and ripping with dissection forceps to mechanically isolate pre-antral follicles from the ovarian cortex in L15 media. Follicles were then fixed and processed with the same protocols as for mouse follicles.

### Microscopy

Samples were imaged on a Leica TCS Sp8 inverted laser-scanning confocal with a white light laser with 2 PMT detectors and 2 HyD detectors using a HCX PL Apo Oil 100x objective, 2048x 2048 pixel resolution, and 0.282 z-step size. Superresolution images were acquired on the Nikon AX NSPARC platform with an Apo TIRF 60x Oil DIC N2 objective, 2.86x zoom, 2048×2048 pixel resolution, and 0.172 z-step size. Images were formatted and cropped with Imaris Software version 10.1.1 (Bitplane).

### Follicle enzymatic digestion and reaggregation

P12-P15 mice were euthanized via cervical dislocation, ovaries were removed and placed in pre-warmed L15 media (11415-064) on a 37C heated microscope stage mounted on a dissecting microscope in a biosafety cabinet with HEPA filter. 2-3 ovaries were placed in 1mL L15 with 25 ug/mL Liberase (cat 5401119001) and 200 ug/mL DNase I (cat 79254) and ripped apart with dissection forceps (cat 11251-20). After 20 minutes a p1000 pipette pre-coated with 30% BSA (cat A30075-100.0) and then “rinsed” 5x in L15 was used to triturate samples for 30 seconds. After another 10 minutes (total of 30) 100 ul of 30% BSA was added and the samples were triturated again. Samples were then added to a 200 um filter (cat 43-10200-40) stacked on a 40 um filter (cat 43-10040-40) and then rinsed. The 200 um filter removes un-dissociated oocyte tissue and the 40 um filter allows primordial and small primary follicles to pass through. The 200 um filter was removed and the 40 um filter was rinsed 3x with L15. The 40 um filter was then inverted and follicles were eluted off with 1 mL 0.05% Trypsin (cat 25300-054). After 10 minutes 100 ul BSA was added and the sample was triturated until all oocytes were bare (about 20 times). The sample was then passed through a pre-rinsed 20 um filter (cat 43-10020-40) to collect granulosa cells. Granulosa cell suspensions were visually inspected to confirm that they contained no oocytes. The 20 um filter was then rinsed 3x with L15 and then inverted and oocytes were eluted off with L15. Oocytes and granulosa cell suspensions were transferred to separate tubes and were then spun at 750 rcf for 5 minutes. As much supernatant was removed as possible without disrupting the cells and replaced with adKSOM (cat MR-101-D) supplemented with 10 nm Estradiol (cat E8875) 100 ng/mL GDF9 (cat 739G9010) and 10 IU/mL FSH (cat F4021).

Oocyte and granulosa cell suspension were then combined according to experimental setup, added to Ultra Low attachment wells (cat 7007) and spun at 750 rcf for 30 seconds. Plates were placed in incubators and removed each of the first 3 days to be spun at 750 rcf for 30 seconds. Half of the supplemented adKSOM was exchanged with fresh supplemented adKSOM on the third 3 day and then left undisturbed in the incubator for 3 more days. Reaggregates were then either fixed with 2% PFA directly in the culture wells for 10 minutes for staining or oocytes were re-isolated.

Oocytes were re-isolated by transferring reaggregates to warm 0.05% Trypsin, with BSA quenching after 5 minutes. Oocytes then were transferred through 3 L15 washes and visualized on a brightfield microscope. Only bare oocytes where granulosa cells were completely removed were selected for RNA isolation.

### RNA extraction

25-35 oocytes were transferred to phase lock gel heavy tubes (cat 2302830). 100 ul of Trizol (cat 15596026) and then 50 ul chloroform (cat 383770010) were added. Tubes were inverted for 15 seconds and then incubated at room temperature for 3 minutes. Tubes were then centrifuged at 12,000 rcf at 4°C for 30 minutes. The aqueous phase was then transferred to a new RNase-free tube and 50 ul ice-cold isopropanol, 10 ul 3M sodium acetate (cat AM9740), and 2.5 ul glycogen (cat R0551) were added. The samples were mixed gently but thoroughly and then placed at -20 for 60 minutes to encourage precipitation. Tubes were then centrifuged at 12,000 rcf at 4°C for 30 minutes. Supernatants were discarded and pellets were rinsed in 150 ul 70% ice-cold ethanol (diluted with RNase-free water) and then centrifuged at 7,500 rcf for 5 minutes at room temperature. This step was repeated. As much ethanol as possible was then removed without disturbing the pellet and tubes were left open under a sterile flame to dry for 5 minutes. Pellets were then resuspended in 10 ul nuclease-free water (cat AM9937) at 65°C for 5 minutes. These samples were then either immediately shipped for sequencing or stored at -80°C.

### EU incorporation and processing

P12-P15 mice were injected with 1 mg 5-ethynyl-uridine (cat C10330 or CCT-1261) dissolved in sterile saline (cat S5815). These mice were sacrificed 16-18 hours later and follicles were either mechanically isolated and processed for imaging or enzymatically isolated for reaggregations. Enzymatically isolated EU-follicles were processed as described above to isolate EU-labelled granulosa cells and re-aggregated with oocytes from a separate litter. Fixed whole follicles were fixed and washed as described above and processed according to Click-iT kit instructions (cat CCT-1261).

### Bioinformatics

Following C57BL/6J (B6) oocytes aggregate with CAST/EiJ (CAST) granulosa cells and subsequent removal of the granulosa cells, total RNA was isolated from three replicates of B6 oocytes as described above. Total RNA samples were sent on dry ice to Novogene, and sequencing libraries were constructed using the low input SMART Seq V4 kit (Takara Bio) according to manufacturer’s protocols. Libraries were sequenced 2 × 150bp paired-end on the NovaSeq platform (Illumina). Approximately 100 million reads were generated in the first sample, and ∼65 million reads were generated from the subsequent two replicates. FastQC^48^ was used to assess read quality, and TrimGalore^49^ was used to remove Illumina adapter sequences and trim the 2 × 150bp reads down to the highest quality 75bp. The forward read of each trimmed pair was used for transcript quantification. Trimmed reads were first aligned to the B6 transcriptome (Genome version: GRCM39. Gene annotation version: v105) using Bowtie aligner^51^, allowing for zero mismatches, with parameters “-best -strata -a -v 0 -un”. Any reads failing to map perfectly to the reference B6 transcriptome were aligned a second time to a “CAST-specific” transcriptome containing all CAST SNPs and indels that differ from the reference. Reads that did not map (or mapped with >= 1 mismatch) to the B6 reference transcriptome but mapped perfectly (0 mismatches) to the CAST transcriptome were considered likely to be of CAST strain origin. Many additional reads of CAST origin will likely be missed by this conservative read trimming and alignment strategy, which was selected to minimize false positive CAST read assignments due to sequencing errors or overly permissive alignment parameters.

To estimate transcript abundance levels from reads of CAST origin, we applied an Expectation-Maximization algorithm as implemented in RSEM (RNA-Seq by Expectation-Maximization)^53^. Gene ontology enrichment analysis was then performed using the *enrichGO* function implemented in the *clusterProfiler* Bioconductor R package^21^ to determine the biological relevance and specific pathways involved in the overlap genes. The STRING biological database was used to determine functional protein association networks^20^.

## Acknowledgements

This work was supported by NIH grants 1R01GM122902 (DJL), 1R01ES023297 (DJL), the Global Consortium for Reproductive Health through the Bia-Echo Foundation GCRLE-0123 (DJL) and and the W.M. Keck Foundation (DJL). C.A.D. was supported by National Institutes of Health training grant T32 HD 007263 and The Jane Coffin Childs Memorial Fund for Medical Research. This publication was made possible with Pilot Program support from the Southwest National Primate Research Center, grant number P51 OD011133.5. S.C.M. was supported by NIH grant R35GM133495. A.T. was supported by the JAX Paigen Fellowship.

Thanks to Kyle Marchuk, Thomas Kenney, and Kyle Wycoff for providing access beyond institutional demos to the NSPARC platform, to James Gardner, Juan Du, Angelica Olmo-Fontanez, Kimberley Phillips, Debbie Christian, Sharon Price, Donna Layne-Colon, Celina Valdez and Samuel Galindo for generous assistance procuring ovary tissues. We thank Liz Gavis, Sandra Schmidt, and all members of the Laird lab for critical feedback on this work.

## Author Contributions

Conceptualization, C.A.D. and D.J.L.; Methodology C.A.D. and D.J.L; Software A.T. and S.C.M; Validation C.A.D.; Formal analysis, all authors; Investigation C.A.D.; Resources D.J.L.; Data Curation A.T. and S.C.M; Writing – original draft, C.A.D. and D.J.L.; Writing – review and editing, all authors; Visualization C.A.D. and A.T.; Supervision D.J.L.; Project administration D.J.L.; Funding Acquisition D.J.L.

## Supplementary Figures and Tables

**SFigure 1.**
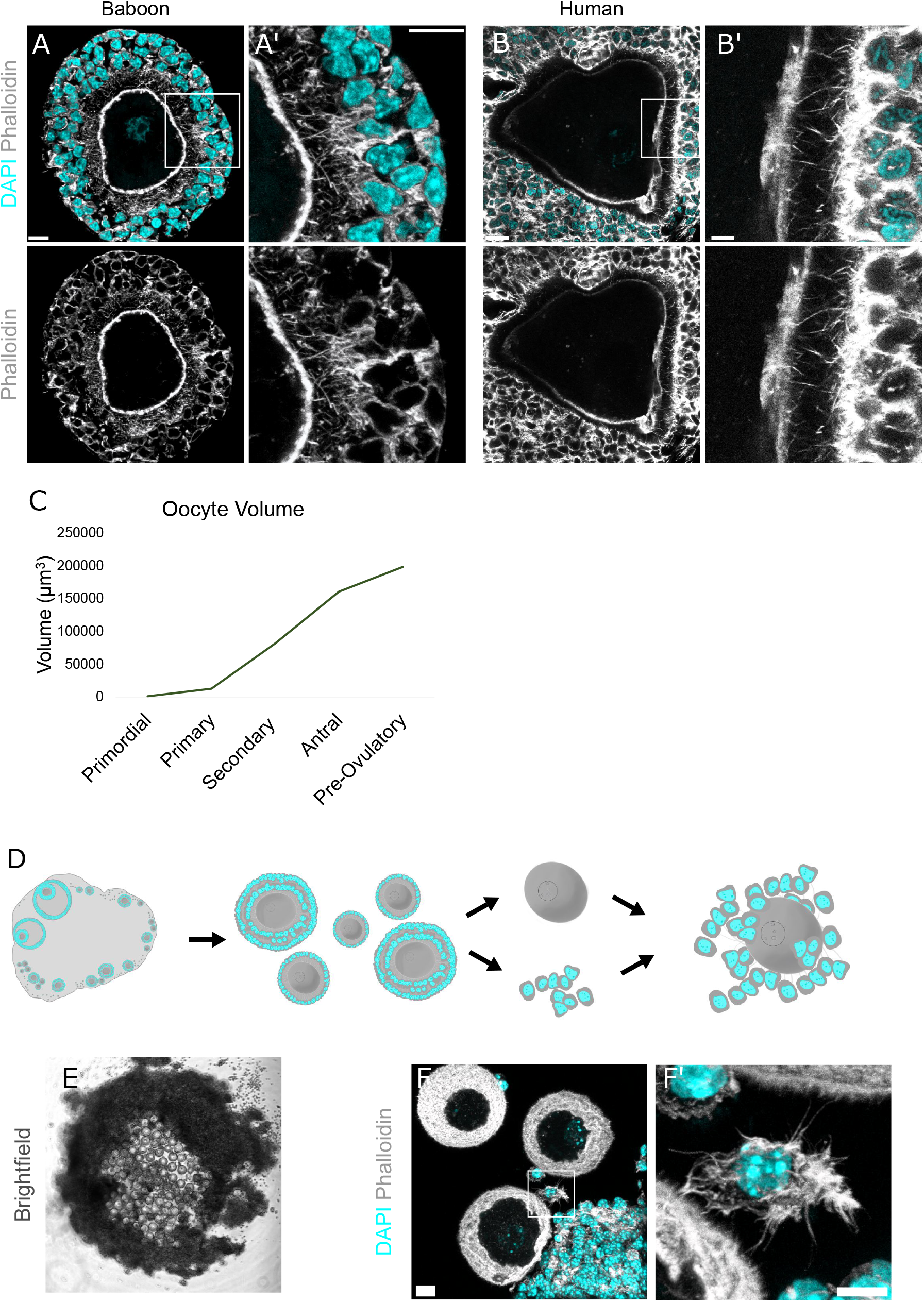
TZPs are observed in Primate preantral follicles and in follicle reaggregations. A. Confocal z-projection of a secondary Anubis baboon follicle, with DNA marked by DAPI (turquoise) and TZPs marked with phalloiding (grey). Scale bars = 10 um (A & A’). B. Confocal z-projection of a secondary human follicle similarly stained to A. Scale bars = 20 um (B), 5 um (B’). C. Mouse oocytes undergo the largest increase in volume between primary and antral stages. Data adapted from (Griffin et al. 2006) D. Schematic of follicle reaggreagation protocol. E. Brightfield image of granulosa cell-oocyte reaggregation in culture. F. Confocal z-projection of granulosa cells that have re-grown TZPs that contact oocytes. Scale bar = 10 um.

**Table 1. Complete list of Castaneous transcripts identified as transferred to B6 oocytes**

**Table 2. Top 50 RNA-Binding Protein (RBP) motif instances in transcripts present in all 3 replicates**

## Notes

### Competing Interest Statement

D.J.L. is on the SAB of Vitra, Inc.

### Summary of Updates

The author order has been corrected in the Biorxiv listing.

